# Coherent scene context accelerates and reshapes neural object representations

**DOI:** 10.64898/2026.06.22.733573

**Authors:** Alireza Javadi, Hamid Soltanian-Zadeh, Karim Rajaei

## Abstract

Coherent scenes facilitate object recognition, but the representational basis of this facilitation and its temporal evolution in the brain remain unclear. We tested this question using EEG and multivariate pattern analysis while 15 participants categorized objects from five semantic categories after a 500-ms preview of either an intact rendered scene or a phase-scrambled version of the same background. Reliable object decoding emerged earlier in intact scenes than scrambled scenes (142 ± 5 vs. 162 ± 10 ms), with higher decoding for intact scenes from 124 to 268 ms after object onset. Cross-condition decoding object information that generalized across scene formats, whereas subtracting cross-condition from within-condition decoding identified an earlier and stronger context-dependent component when scene structure was coherent. Cross-temporal representational similarity analysis (RSA) further showed that representational structure established during late scene preview generalized to early object processing only for intact scenes, linking contextual facilitation to anticipatory scene-derived representations. Finally, model-to-brain RSA showed that a language-aligned model explained neural representational geometry in intact scenes better than vision-only models, an advantage attenuated by scene scrambling. These findings indicate that coherent scene context shapes object coding by accelerating object-selective processing and contributing context-dependent representational structure beyond a context-invariant object code.

## Introduction

Objects are typically encountered within structured scenes that provide strong statistical regularities about which objects are likely to occur and where they are likely to appear (Bar, 2004; Oliva and Torralba, 2007). Such regularities provide semantic and spatial priors that can constrain object perception. Behaviorally, these priors facilitate object recognition, particularly when bottom-up object information is degraded or ambiguous (Davenport and Potter, 2004; Munneke et al., 2013). Neural studies have linked such benefits to interactions between scene-selective and object-selective systems, whereby scene context sharpens category-specific object representation in visual cortex (Bar, 2004; Brandman and Peelen, 2017; Wischnewski and Peelen, 2021; Leticevscaia et al., 2024; Soltandoost et al., 2025). However, the extent to which coherent scene context alters the time course and representational content of object processing remains unclear.

One unresolved issue is what scene context adds to neural object representation. Contextual facilitation could reflect an earlier or stronger expression of an object code that is largely invariant across backgrounds. Alternatively, coherent scenes may contribute additional representational structure tied to the semantic and spatial relations between an object and its surrounding scene. These alternatives are not mutually exclusive, but they make different predictions: the first predicts information that generalizes across intact and disrupted scene formats, whereas the second predicts a context-dependent component that is selectively expressed when coherent scene structure is preserved.

A related issue concerns timing. In everyday vision, global scene information can be available before focal object processing is complete. Such prior scene information may establish expectations about likely objects and thereby constrain subsequent object processing (Bar, 2004; De Lange et al., 2018). If contextual facilitation has an anticipatory component, the representational structure elicited during scene preview should be related to the earliest stages of object-evoked activity. Testing this relationship can clarify whether context acts only after object onset or whether scene-derived structure is already in place before the object appears.

A final question is which computational representations best capture context-sensitive neural object code. Deep neural networks (DNNs) provide explicit representational spaces that can be compared with neural data using representational similarity analysis (RSA). Vision-only models may capture image-based similarity, whereas vision–language models trained on image–text pairs may better capture semantic and relational structure relevant to objects embedded in meaningful scenes (Wang et al., 2022; Conwell et al., 2024; Doerig et al., 2025; Rajaei et al., 2026). Thus, if coherent scenes add semantically structured information to object representations, language-aligned models should show stronger alignment with neural representations in intact scenes than when scene structure is disrupted. Here, we combined EEG with multivariate decoding and representational similarity analysis while participants viewed a 500 ms scene preview followed by a briefly presented object embedded either in an intact rendered scene or in a phase-scrambled version of the same scene. This design allowed us to ask whether coherent context accelerates or strengthens object-discriminative information, whether it contributes a context-dependent representational component beyond context-invariant object code, and whether scene-preview representations are reinstated during early object processing. We then compared neural representational geometry with supervised vision, self-supervised vision, and language-aligned vision–language models to test which computational representations best captured object coding when coherent scene structure was available.

### Experiments and Results

To determine how scene context shapes object processing over time, we organized the Results into four complementary analyses. The first three focus on the human EEG data, and the fourth compares those neural representations with computational models. We begin with time-resolved within-condition decoding in intact and scrambled scenes to establish when object-selective information emerges and whether coherent scene structure accelerates and strengthens object processing. We then use cross-condition decoding, together with temporal generalization, to distinguish object representations that are shared across scene contexts from those that depend specifically on coherent scenes, and to assess how stable these representations remain over time. Next, leveraging the scene-preview period that preceded object onset, we apply cross-temporal representational similarity analysis to test whether representational structure formed during scene viewing is reinstated during early object encoding, consistent with an anticipatory influence of scene context. Finally, we compare the EEG representational geometry with that of three deep neural network models trained under different supervisory regimes (supervised vision, self-supervised vision, and language-aligned vision-language models) to ask which computational framework best captures object representations when coherent scene structure is available.

### Coherent scene context accelerates and enhances object decoding

We tested whether coherent scene context facilitates neural object processing and, critically, whether such facilitation reflects changes in the representational structure of object information rather than a nonspecific increase in signal strength. Fifteen participants viewed 25 objects from five semantic categories (food, electronics, containers, office supplies, and home décor) while EEG was recorded (Fig. 1A; full set in Fig. S1). Objects appeared under two context conditions: an intact condition, in which objects were embedded in coherent scenes, and a scrambled condition, in which the same object was presented at the same location on a phase-scrambled version of the same scene background (Fig. 1B). On each trial, participants viewed the scene background for 500 ms, followed by an 80-ms presentation of the target object embedded in the scene (Fig. 1C), and then categorized the object in a five-alternative choice task. Each object was presented up to 30 times per condition. Categorization accuracy was high in both conditions, with a small advantage for intact over scrambled scenes (97.1 ± 0.8% vs. 95.7 ± 1.1%, mean ± SEM; intact > scrambled in 11 of 15 participants; two-tailed Wilcoxon signed-rank test, p = 0.041). Response rate did not differ between conditions (response rate 95.8% vs. 96.3%; p = 0.49), indicating comparable task engagement regardless of scene structure.

**Figure 1.**
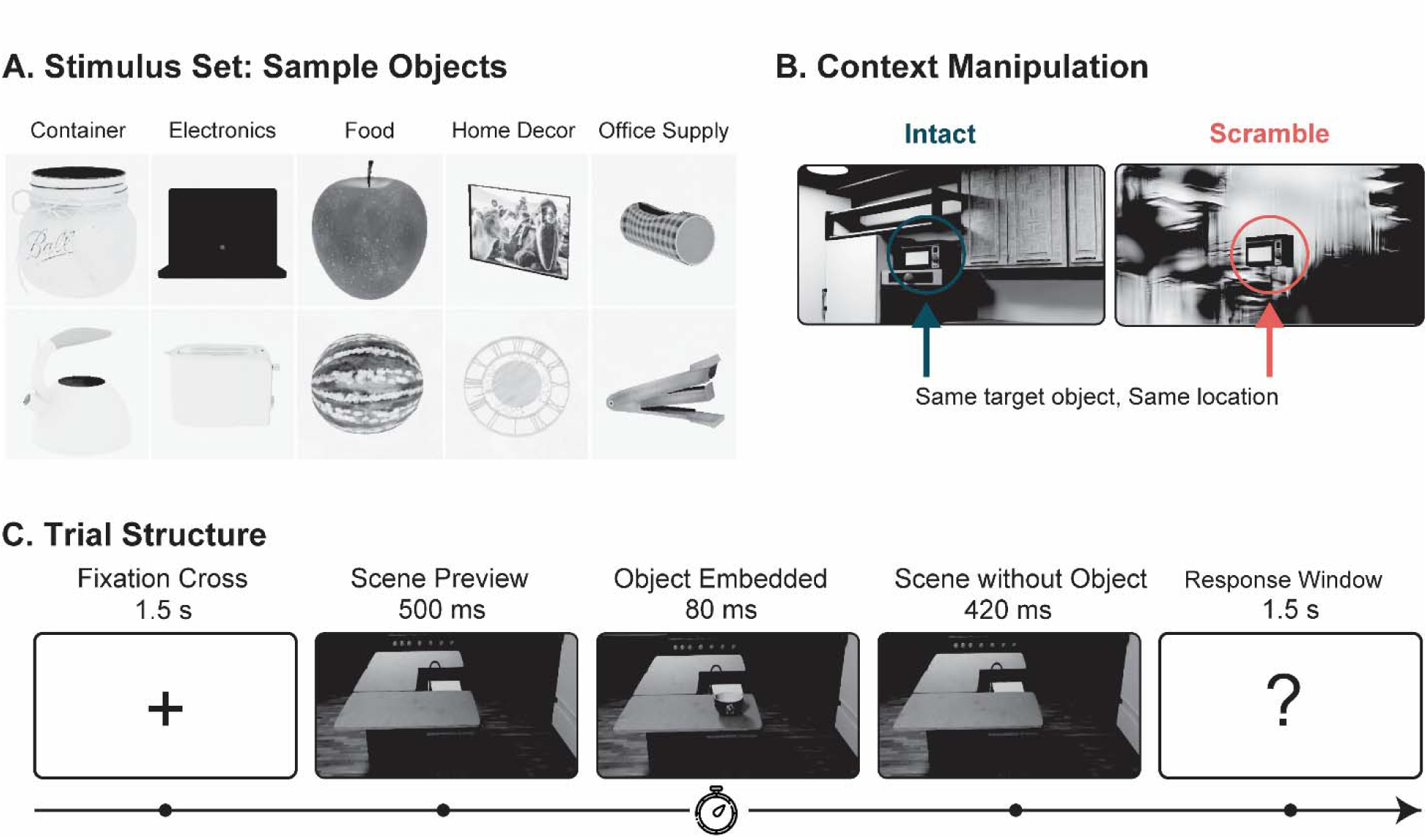
Experimental design. **A.** Example target objects drawn from five semantic categories: containers, electronics, food, home décor, and office supplies. All stimuli were presented in grayscale. **B.** Two context conditions. Target objects were presented on either intact or phase-scrambled scene backgrounds. **C.** Trial structure. Each trial began with a 1.5-s fixation period, followed by a 500-ms preview of the scene background. The target object was then embedded in the scene for 80 ms, after which the background scene alone remained on the screen for 420 ms. Participants categorized the target object by pressing one of five predefined response keys within a 1.5-s response window.

To examine the temporal dynamics of object-selective neural information, we performed time-resolved pairwise decoding separately for the intact and scrambled conditions. At each time point, classifiers discriminated between pairs of objects using multichannel EEG patterns, such that decoding accuracy indexed the discriminability of object representations over time (Fig. 2A).

**Figure 2.**
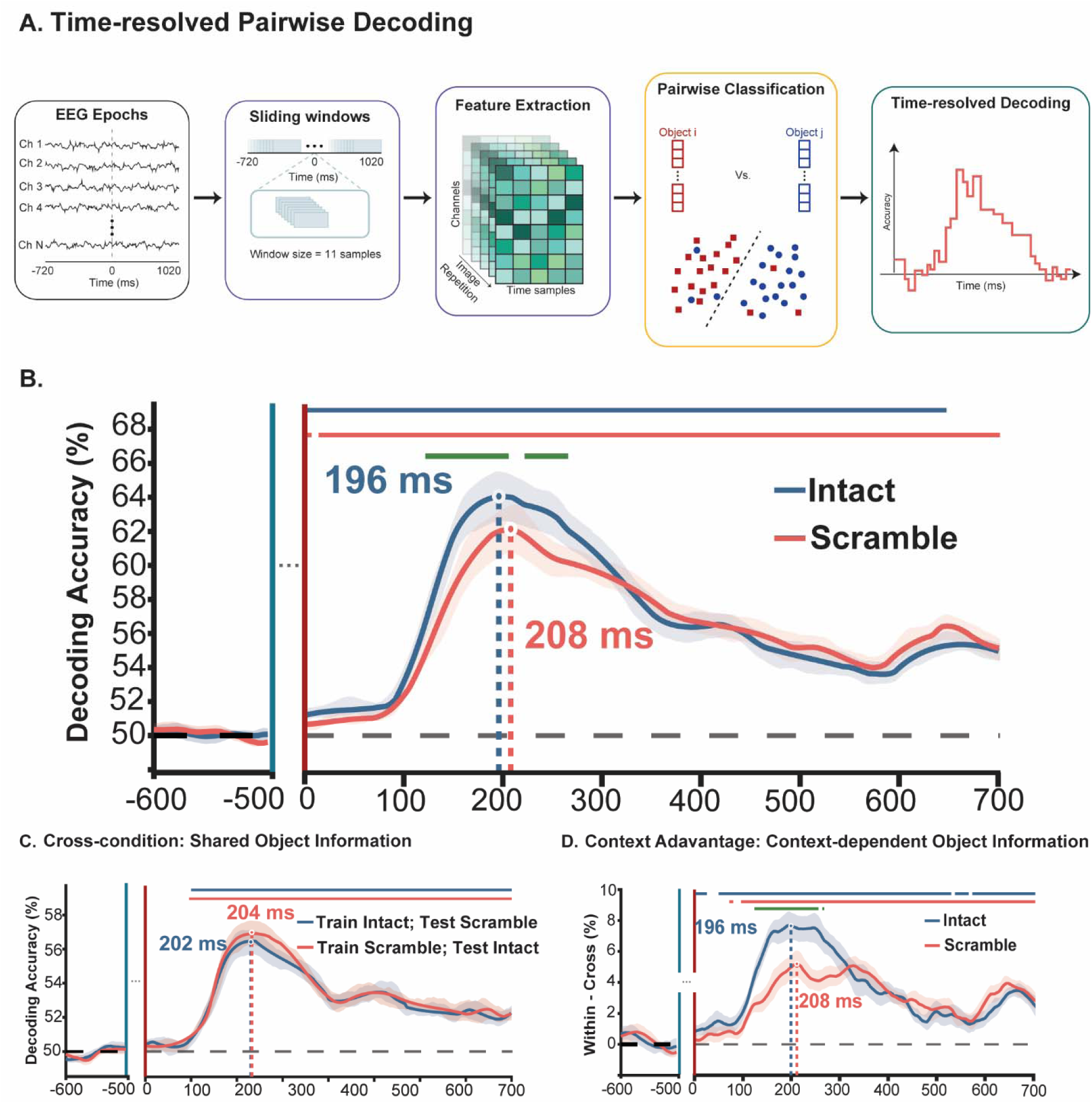
Multivariate pattern decoding results. **A.** Schematic of the time-resolved pairwise decoding analysis. For each participant, EEG epochs were segmented into overlapping sliding windows (11 samples, 44 ms). Within each window, voltages across all retained channels and samples served as features. For every pair of the 25 objects, a linear classifier was trained to discriminate the two objects, and accuracy at each time point yielded a time-resolved decoding curve. **B.** Time-resolved object decoding for intact (blue) and scrambled (pink) conditions. Shaded regions denote ±1 SEM across participants. Horizontal bars indicate time points at which decoding was significantly above chance within each condition. The green bar marks the interval showing a significant difference between conditions (FDR-corrected). Dotted vertical lines indicate peak decoding latency for each condition. **C.** Cross-condition decoding. Time-resolved decoding accuracy for classifiers trained in one scene condition and tested in the other: intact-to-scrambled (blue) and scrambled-to-intact (pink). **D.** Context advantage. The context advantage was defined as within-condition decoding minus cross-condition decoding, indexing scene-dependent contributions to object decoding.

Object decoding emerged earlier in the intact condition than in the scrambled condition. Decoding onset, defined as the earliest time point at which performance rose reliably and remained consistently above chance, occurred significantly earlier for intact scenes (142 ± 5 ms) than for scrambled scenes (162 ± 10 ms; p = 0.014), indicating an approximately 20-ms acceleration in the emergence of reliable object representations when scene context was coherent (Fig. 2B). Scene coherence also affected the timing of peak decoding, which was reached earlier for objects presented intact than in scrambled scenes (196 ± 3.3 ms vs. 208 ± 2.5 ms; p = 0.001).

A direct time-resolved comparison further showed that decoding accuracy was significantly higher in the intact than in the scrambled condition from 124 to 268 ms after object onset (p < 0.05, FDR-corrected). This pattern was consistent across individual participants and across object categories (Fig. S2). Also, representational structure at decoding landmarks is shown as RDMs (Fig. S3A) and their two-dimensional multidimensional scaling (MDS) embeddings (Fig. S4A). Together, these findings indicate that coherent scene context facilitates object processing by accelerating the emergence of object-selective information and increasing its decodability. At the same time, because each object identity was paired with a scene image within each condition, within-condition decoding may reflect both object identity and condition-specific scene information. To dissociate these contributions, we next used cross-condition analyses to isolate object information that generalized across contextual formats.

### Cross-context decoding reveals a shared object code across scene formats

To isolate object-related information from activity associated with the surrounding scene, we performed cross-condition decoding by training classifiers in one context condition and testing them in the other. Information that generalizes across intact and scrambled scenes provides an estimate of object-discriminative information shared across contextual formats.

Cross-condition decoding rose reliably above chance shortly after object onset and showed highly similar time courses in both train-test directions (Fig. 2C). Onset latency did not differ between classifiers trained on intact scenes and tested on scrambled scenes and those trained on scrambled scenes and tested on intact scenes (intact-to-scrambled: 140 ± 2 ms; scrambled-to-intact: 140 ± 2 ms; difference not significant). Peak latency was likewise comparable across train-test directions (intact-to-scrambled: 202 ± 7 ms; scrambled-to-intact: 204 ± 7 ms; difference not significant), and decoding accuracy did not differ significantly at any time point. These results indicate that a substantial portion of object-discriminative information generalized across the two context conditions, consistent with a shared representation of object identity across contextual formats.

### Coherent scene context generates earlier and stronger context-dependent object information

The shared code above cannot, by definition, capture information that depends on the specific scene context in which an object appears. To test our second hypothesis — that coherent context contributes object information beyond this context-invariant code — we isolated the context-dependent component by subtracting cross-condition from within-condition decoding, yielding a context advantage measure. This measure indexes object information that is available within a given contextual format but does not generalize across formats, providing a direct estimate of a context-dependent component of object-related discriminability.

Coherent scene context shaped this context-dependent component in two ways. First, it accelerated its emergence: the context advantage arose approximately 23 ms earlier for intact than for scrambled scenes (128 ± 2 ms vs. 151 ± 4 ms; p < 0.0122). Second, it amplified its magnitude: the context advantage was significantly greater for intact than scrambled scenes from 121 to 256 ms after object onset (p < 0.05, FDR-corrected; Fig. 2D). Thus, beyond the object code shared across formats, coherent scene structure gives rise to a context-dependent component of object representation that is both earlier and stronger, supporting the idea that scene context contributes object information over and above a context-invariant representation.

### Coherent scene context advances object representations to a later processing stage

We next examined the temporal generalization of object representations, in which classifiers were trained at one time point and tested across all other time points. Within-condition temporal-generalization matrices (Fig. 3A, D) revealed two phases of representational dynamics. In the early phase (before ∼400 ms), significant decoding was concentrated near the diagonal, consistent with transient and temporally specific representations. In the later phase, from ∼400 ms onward, decoding generalized more broadly to off-diagonal time-points, indicating a more stable and sustained representational format.

**Figure 3.**
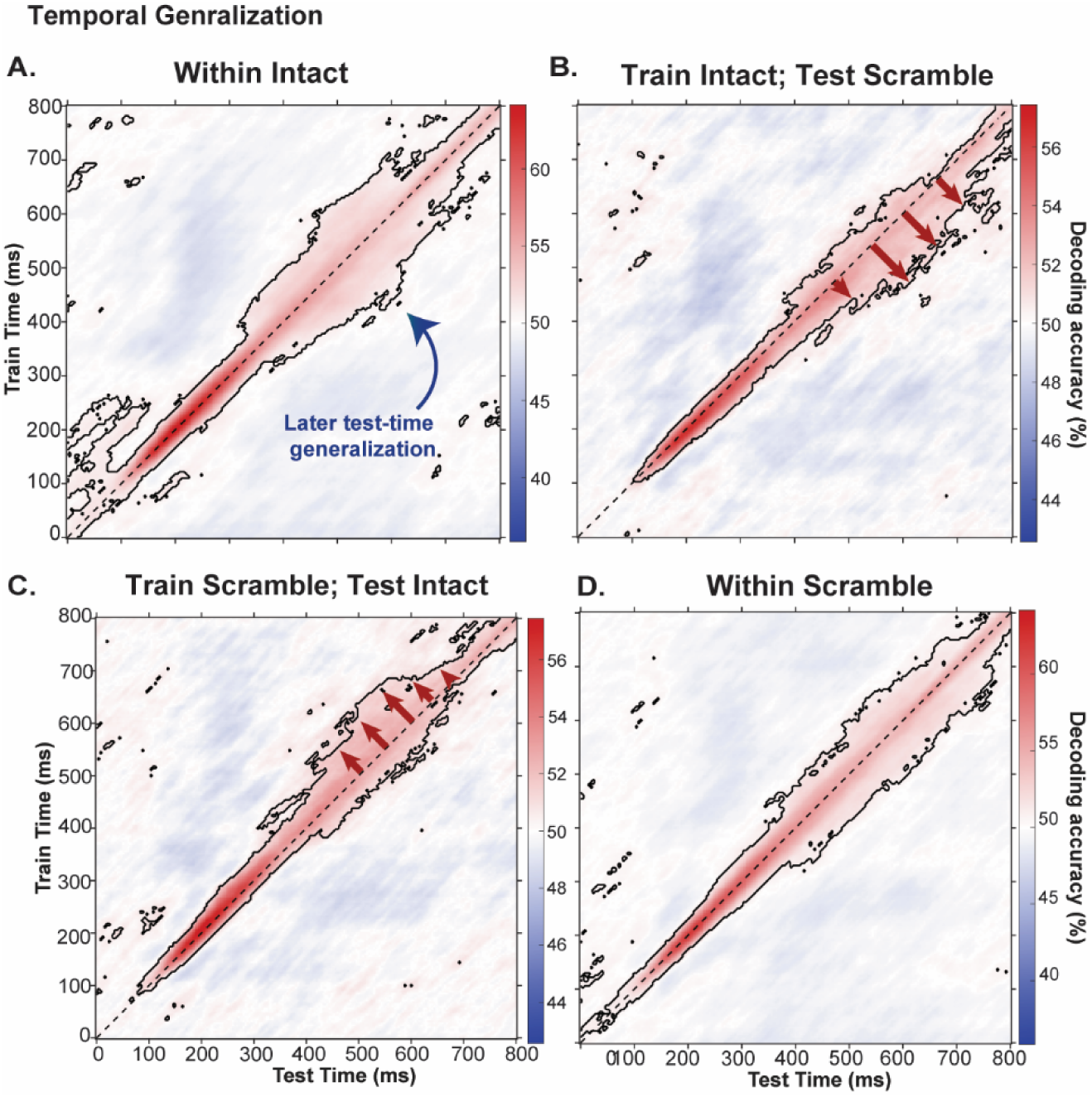
Temporal Generalization (A-D). Within- and cross-condition temporal generalization for intact and scrambled conditions. A classifier was trained on the EEG pattern at each time point and tested on the patterns from all other time points. Color indicates decoding accuracy, and black contours indicate significant clusters (FDR-corrected). Arrows indicate the asymmetry of temporal generalization from the diagonal.

Cross-condition temporal generalization revealed a directional asymmetry that speaks directly to the functional role of coherent context: representations the visual system forms early under coherent scenes are the same ones it reaches only later when scene structure is disrupted (Fig. 3B, C). Specifically, decoding generalized preferentially toward the scrambled condition, such that representations present at earlier time points in the intact condition generalized to later time points in the scrambled condition. This asymmetry was most pronounced at later stages of processing. In other words, coherent scene context advances the object-processing trajectory in time, allowing the system to arrive sooner at representational states that disrupted scenes attain only after additional processing. This benefit, therefore, extends beyond the early gain in decoding accuracy, shaping later stages of object processing associated with higher-level visual and semantic representations.

### Scene preview elicits a representation that generalizes to early object processing

Because our paradigm included a scene-preview period before object onset, we next asked whether viewing the scene alone elicited object-relevant representations that could bias subsequent object processing. This speaks directly to anticipatory accounts of contextual facilitation, which propose that coherent scene context establishes neural representations before object onset that may constrain incoming object processing (Bar, 2004). To test this, we compared the representational geometry of neural activity during the scene-preview period with that elicited after object onset using cross-temporal representational similarity analysis (Fig. 4A).

**Figure 4.**
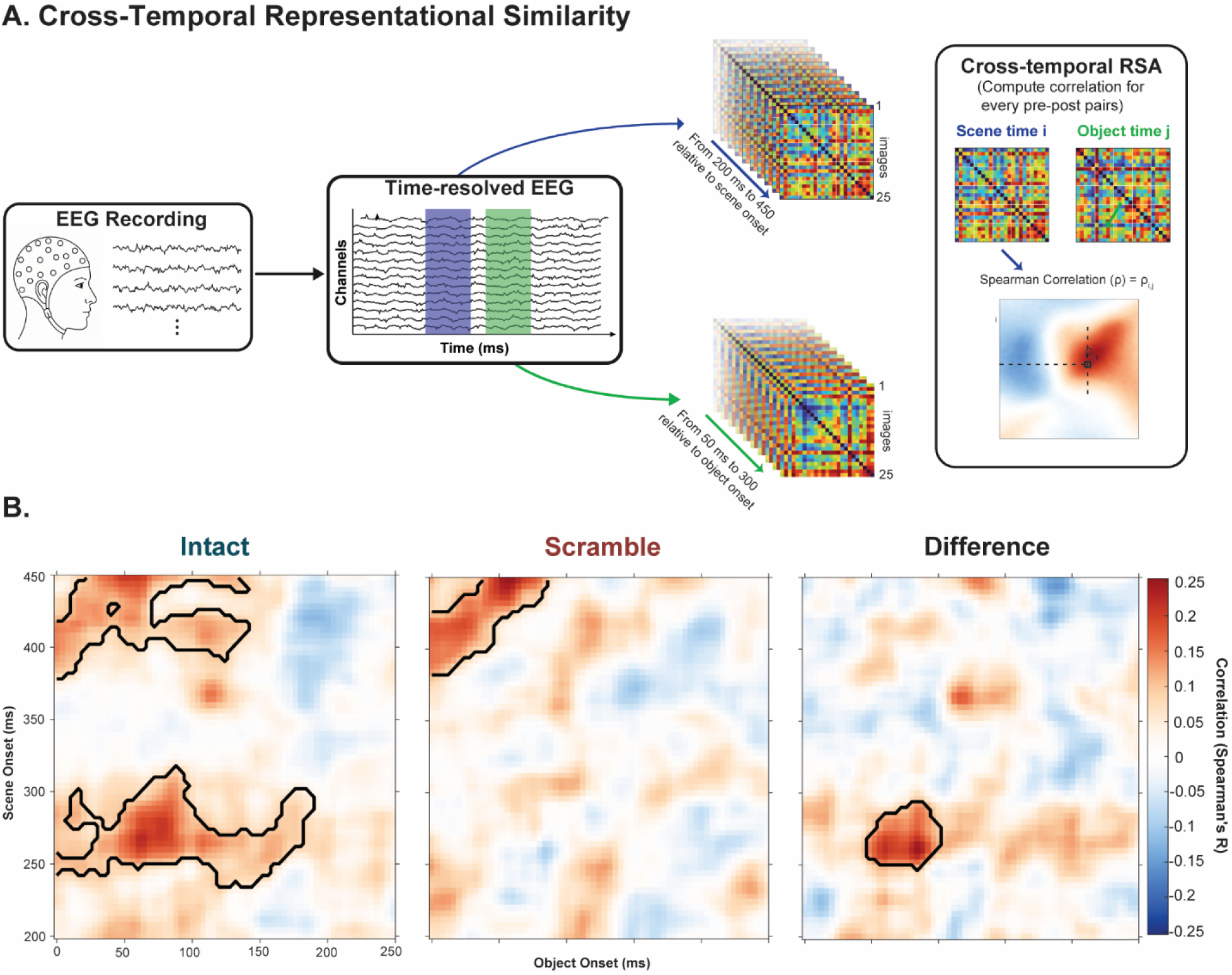
Cross-temporal representational similarity analysis. **A.** Schematic of the cross-temporal representational similarity analysis. Time-resolved 25 × 25 representational dissimilarity matrices (RDMs) were computed from EEG during the scene-preview period (relative to scene onset) and during the post-object period (relative to object onset). For each pair of pre- and post-object time points, the corresponding RDMs were correlated (Spearman’s ρ), yielding a two-dimensional cross-temporal similarity matrix **B.** Cross-temporal representational similarity analysis and resulting similarity matrices for intact scenes, scrambled scenes, and their difference (intact minus scrambled). The vertical axis indicates time relative to scene onset, and the horizontal axis indicates time relative to object onset. Color represents the Spearman correlation between scene-period and object-period RDMs. Black contours indicate significant clusters (cluster permutation test, p < 0.05).

In the intact condition, this analysis revealed significant generalization between late scene processing and early object processing, indicating that the representational geometry established during scene viewing was reinstated during early stages of object processing (Fig. 4B, Intact). This result suggests that scene-derived representations, formed before object onset, influence the processing of the incoming object, consistent with the view that coherent scenes do more than enhance a context-invariant object code but contribute an additional context-dependent representational component. No comparable effect was observed in the scrambled condition, despite matched low-level image statistics, indicating that this generalization depended on coherent semantic and spatial scene structure rather than low-level background statistics alone.

The effect was confined to a specific temporal window: neural patterns from 250–300 ms after scene onset generalized significantly to patterns observed 50–100 ms after object onset in the intact condition (Fig. 4B, Difference). Notably, this 50–100 ms post-object-onset window precedes the mean decoding onset we observed in the scrambled condition (162 ± 10 ms), and overlaps with the accelerated decoding onset in the intact condition (142 ± 5 ms), suggesting that the reinstated scene-preview structure may contribute to this earlier emergence of object-discriminative information. Together, these findings indicate that coherent scene preview establishes representational structure that is linked to — and may facilitate — the accelerated emergence of object representations after stimulus onset, consistent with the anticipatory framework motivating our experimental design.

### Language-aligned model representations show better alignment in coherent scenes

To examine which computational representations best captured the context-sensitive object representation observed in the EEG data, we compared neural representational geometry with that of three deep neural network models trained under different supervisory regimes: ViT-ImageNet, a supervised vision-only model trained on image labels (Dosovitskiy et al., 2020); DINOv2, a self-supervised vision-only model trained without labels (Oquab et al., 2023); and MetaCLIP, a vision–language model trained on image–text pairs (Xu et al., 2023). For each model and time point, we constructed a 25 × 25 representational dissimilarity matrix (RDM) over the object set. Model-brain alignment was then quantified as the variance explained (R²) between model and EEG RDMs, separately for intact and scrambled conditions (Fig. 5A). This analysis was motivated by the possibility that, if coherent scene context introduces a context-dependent representational component—beyond a context-invariant object code — then models encoding semantic and relational structure should show preferentially stronger alignment when scene coherence is preserved.

**Figure 5.**
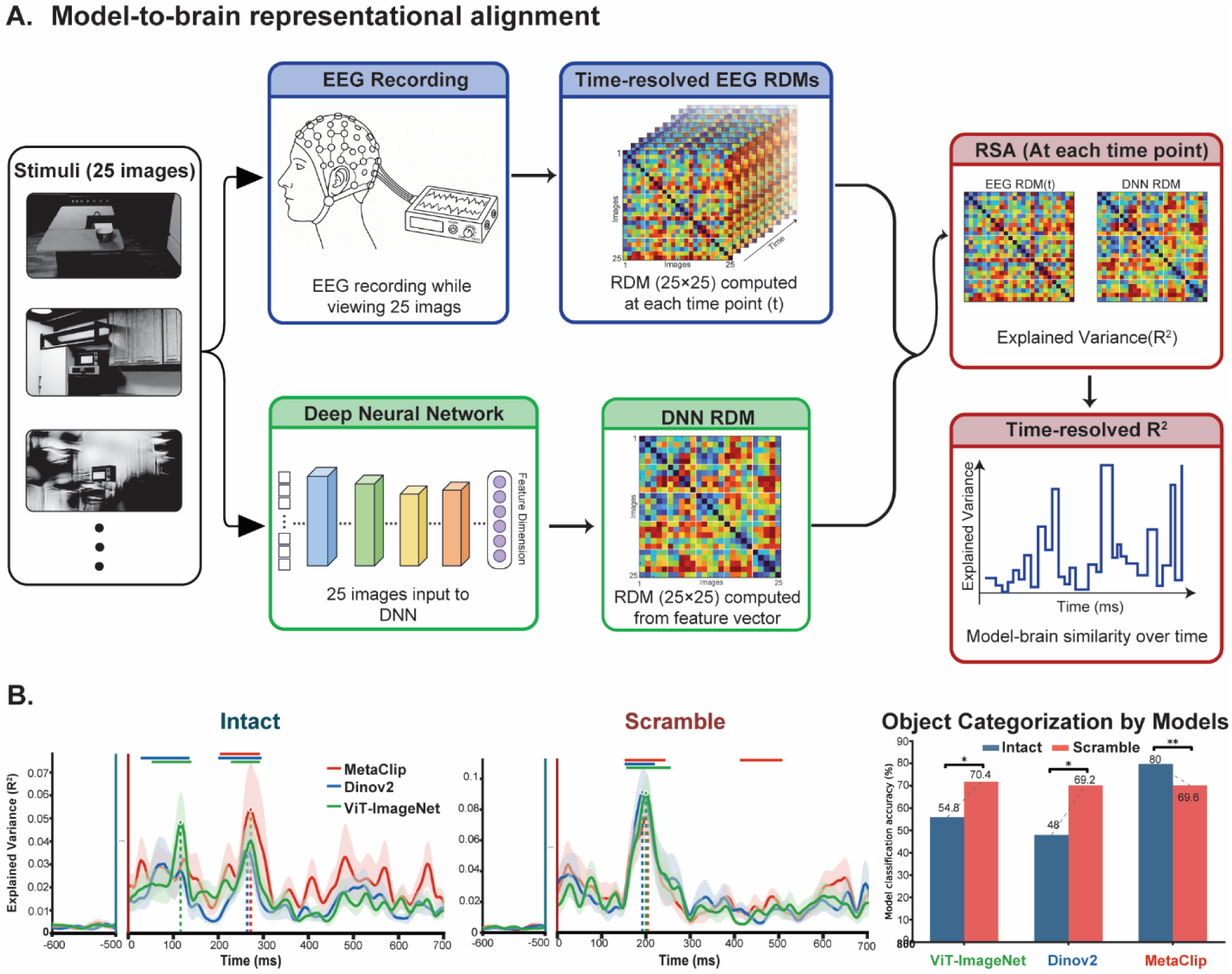
Model-to-brain representational alignment. **A.** Schematic of the model-to-brain representational alignment analysis. The same 25 object images shown to participants were passed through each deep neural network, and a 25 × 25 RDM was computed from the resulting feature vectors. At each time point, the model RDM was correlated with the EEG RDM (Spearman’s ρ) and converted to explained variance (R²), giving a time-resolved alignment curve per model and condition. **B.** Model-to-brain representational alignment. Time-resolved explained variance (R²) between brain 25 × 25 representational dissimilarity matrix (RDM) and model RDM in the intact and scrambled conditions. Results are shown for MetaCLIP (a vision–language model trained on image–text pairs; red), DINOv2 (a self-supervised vision model; blue), and ViT-ImageNet (a supervised vision model; green). Shaded regions denote ±1 SEM (standard error of the mean) across participants. Horizontal bars indicate time points showing significant model–brain alignment for each model (cluster-corrected, p < 0.05), and dotted vertical lines mark peak latency for each model. Performance of object categorization across models. Blue bars represent the intact condition, and pink bars represent the scrambled condition. * p < 0.05, ** p < 0.01.

All three models showed significant alignment with neural object representations in both conditions, but their temporal profiles diverged as a function of scene coherence. In intact scenes, the two vision-only models aligned with the EEG data at relatively early latencies (ViT-ImageNet: 101 ± 2 ms; DINOv2: 64 ± 50 ms), whereas MetaCLIP reached significance later (222 ± 81 ms). Peak latencies followed a similar ordering: ViT-ImageNet peaked earliest, at 116 ms (peak R² = 0.048 ± 0.013), while DINOv2 and MetaCLIP peaked later, at 264 ms (peak R² = 0.037 ± 0.010) and 272 ms (peak R² = 0.054 ± 0.021), respectively (EEG and model RDMs at peak in Fig. S3B; MDS in Fig. S4B). Critically, MetaCLIP explained significantly more variance at its peak than either ViT-ImageNet or DINOv2 (both p < 0.05), indicating stronger late correspondence with neural representational geometry when scene structure was coherent.

In scrambled scenes, this pattern was largely absent. Alignment latencies were closely matched across models (ViT-ImageNet: 174 ± 3 ms; DINOv2: 172 ± 0 ms; MetaCLIP: 164 ± 2 ms), and all three reached peak alignment within a comparable time window (ViT-ImageNet: 200 ms; DINOv2: 192 ms; MetaCLIP: 204 ms), with no significant pairwise differences in peak latency or peak R² (Fig. 5B Scramble). The selective emergence of MetaCLIP’s advantage under intact scenes — paralleling the earlier decoding onset (142 ± 5 ms vs. 162 ± 10 ms) and the scene-preview generalization effect reported above—suggests that the additional representational component conferred by coherent context is specifically captured by language-aligned features.

Model classification performance was consistent with this interpretation. Only MetaCLIP showed a contextual advantage, classifying objects more accurately in intact than scrambled scenes (80.0% vs. 69.6%; p = 0.0007). Both vision-only models performed worse in intact than scrambled scenes (ViT-ImageNet: 54.8% vs. 70.4%; p < 0.0003; DINOv2: 48.0% vs. 69.2%; p < 0.0001; Fig. 5B), suggesting that coherent scene backgrounds disrupted vision-only categorization while providing a representational benefit to the language-aligned model. Together, these findings indicate that coherent scene structure selectively strengthens model–brain alignment for a vision–language model, consistent with the proposal that semantic and relational scene regularities contribute a context-dependent representational component to object coding that is captured more fully by models encoding language-grounded semantic structure. This pattern held across multiple architectures within each supervisory family (Fig. S5).

## Discussion

The present study examined whether coherent scene context influences how the brain represents objects. Our findings suggest that object representations comprise two separable components: a context-invariant object code that generalizes across contexts, and a context-dependent representational component that is selectively present when scene structure is preserved. The object-related component likely reflects processing that is relatively independent of background structure, whereas the context-related component appears to provide additional representational information tied to the object’s embedding within a coherent scene, and was largely absent when scene structure was disrupted. Rather than simply making discriminative object information available earlier or more strongly, coherent scenes thus contributed an additional representational component—a distinction directly motivating the two hypotheses set out in the Introduction. This contextual contribution increased object discriminability and accelerates the emergence of decodable object representations, advancing decoding onset from 162 ± 10 ms to 142 ± 5 ms.

One plausible mechanism underlying this facilitation is object-related expectation. Top-down accounts propose that coherent scenes can pre-activate likely object representations before the object appears, thereby biasing subsequent object processing (Bar, 2004; Brandman and Peelen, 2017; De Lange et al., 2018; Wischnewski and Peelen, 2021; Krugliak et al., 2024). The scene-preview design used here may have favored such a mechanism because it allows scene representations to develop before object onset. This contrasts with earlier MEG studies in which scenes and objects were presented simultaneously and contextual facilitation of object representations emerged relatively late, approximately 320 ms after stimulus onset (Brandman and Peelen, 2017; Leticevscaia et al., 2024). In the present paradigm, the context advantage, measured as the intact-minus-scrambled difference in decoding accuracy, emerges approximately 120 ms and peaks around 200 ms after object onset, roughly 100 ms earlier than in simultaneous-presentation paradigms. We interpret this as evidence that, when scene context is available before object onset, anticipatory processing can shift contextual modulation of object representations to substantially earlier time-points relative to object presentation. Consistent with this interpretation, cross-temporal RSA shows that representational geometry established during late scene preview re-emerged during early object processing, suggesting that scene-derived information shapes the earliest stages of object-evoked activity. The absence of a comparable effect in the scrambled condition, despite preserved low-level image statistics, further indicates that this effect depends on the semantic and structural coherence of the scene rather than on low-level visual properties alone. It is worth noting that these two observations are not contradictory: the temporal generalization analysis tracks the magnitude of the context advantage, which accumulates across time and is most pronounced at later processing stages; the cross-temporal RSA, by contrast, tracks the origin of the context-dependent representational geometry, showing that it is established during scene preview and re-expressed from the earliest latencies of object-evoked activity onward.

The model comparison further suggests that the structure of neural object representations in coherent scenes is better captured by language-aligned than by vision-only models. The language-aligned model (MetaCLIP) showed a relative alignment advantage in intact scenes that was specific to later processing stages, whereas no comparable advantage is observed in scrambled scenes. This pattern is consistent with the possibility that language supervision supports representations that better capture the semantic and relational regularities linking objects to their scene context—precisely the regularities that the scrambling manipulation removes. By contrast, vision-only models may capture visual similarity well but be less sensitive to the broader contextual structure that organizes objects in natural environments. Notably, the two vision-only models classify objects more accurately when backgrounds are phase-scrambled than when they are intact, the opposite of the human pattern. This suggests that, in the absence of semantic supervision, coherent background features compete with target object features rather than supporting their interpretation.

These results converge with previous findings that language-aligned models capture brain representations of natural scenes better than vision-only models (Wang et al., 2022; Doerig et al., 2025), and with behavioral evidence also shows that language-aligned models capture human-like contextual facilitation in object recognition, whereas vision-only models—supervised or self-supervised—do not (Rajaei et al., 2026).

Several limitations should be noted. First, the present paradigm used a single difficulty level in which objects were clearly visible. Prior work has shown that contextual facilitation increases with object recognition difficulty, with stronger effects for degraded, occluded, or non-canonically presented objects (Rajaei et al., 2026). How the representational pattern reported here generalizes to more challenging viewing conditions remains to be tested. Second, each object identity was consistently paired with a specific background in the intact condition. Although the cross-condition analyses help isolate context-invariant object information, within-condition decoding and context-advantage measures could still reflect a mixture of object identity, background identity, and object-background association. Third, the stimulus set was restricted to indoor scenes and indoor objects. Outdoor environments exhibit distinct spatial and semantic regularities, and whether the present findings extend to these contexts remains an open question.

## Materials and Methods

### Participants

Fifteen volunteers (10 male; mean age 27.27 years, SD 4.54) with normal or corrected-to-normal vision participated in the study. All gave written informed consent. The protocol was approved by the Ethics Committee of Tarbiat Modares University (approval ID: IR.MODARES.REC.1404.083).

### Scene–object Stimulus Set

To manipulate scene–object relationships under controlled conditions while preserving naturalistic structure, stimuli were rendered using the OmniGibson embodied-AI environment (Ge et al., 2024; Li et al., 2024), which supports fine-grained control over scene layout, viewpoint, and lighting. Twenty-five object identities were drawn from five high-level semantic categories—food, electronics, containers, office supplies, and home décor (five objects per category)—defined using metadata from the THINGSplus dataset (Stoinski et al., 2023).

Each object was placed within a semantically appropriate scene (e.g., microwaves in kitchens, folders in workspaces, apples on tables) using OmniGibson’s affordance-aware scripting tools, and placements were manually verified for ecological plausibility (Rajaei et al., 2026). To minimize trivial pixel-level regularities, camera viewpoint and lighting were varied across renders. Each object identity was, however, consistently paired with the same rendered scene image across repetitions in the intact condition. This design increased within-object consistency for time-resolved decoding but also entails that object identity is statistically associated with its background; within-condition decoding can therefore reflect a mixture of object- and background-related information. Our main inferences are based on contrasts that match object identity, object position, and low-level background statistics across conditions (intact vs. scrambled). All images were converted to grayscale to remove color-based saliency. Grayscale was a deliberate choice to remove color as a low-level confound, so condition differences can’t be attributed to chromatic cues.

### Phase-scrambled Images

To disrupt global scene structure while preserving low-level image statistics, we generated phase-scrambled versions of the scene backgrounds. Each background image was transformed into the Fourier domain via FFT; the phase spectrum was randomized while the amplitude spectrum was preserved, and an inverse FFT yielded the scrambled image. Phase scrambling was applied to the background only; the target object was left intact and superimposed at the same location used in the intact condition. Intact and scrambled trials were thus matched on object identity, object position, and amplitude-derived background statistics, while differing in the presence of coherent global scene structure (Portilla and Simoncelli, 2000). (https://github.com/TetsuyaOdaka/texture-synthesis-portilla-simoncelli)

### Task and Procedure

Participants performed an object categorization task in which target objects appeared briefly within either an intact scene or its phase-scrambled counterpart, with the same scene background. Trials from the two conditions were intermixed in random order.

Each trial began with a central red fixation cross on a black background (1500 ms), followed by a 500-ms preview of the scene background alone. The target object was then embedded on the background for 80 ms, after which the background remained on screen for an additional 420 ms. Each object identity appeared approximately 30 times per condition.

To minimize motor-related EEG activity time-locked to object onset, explicit responses were required only on intermittent probe trials (every one to five trials, randomly determined). On probe trials, participants reported the category of the object presented on the immediately preceding trial via a five-alternative keypress (one key per category) within a 1500-ms response window. Because participants did not know in advance which trials would be probed, object categorization remained task-relevant throughout. Stimulus presentation and response collection were controlled in MATLAB with the Psychophysics Toolbox (Brainard, 1997).

### Behavioral analysis

Categorization performance was assessed on trials, on which participants reported the category of the object presented on the preceding trial. For each participant and condition (intact, scrambled), accuracy was defined as the number of correct responses divided by the number of trials on which a response was given (responded-only definition); trials with no response were excluded from the accuracy measure and quantified separately as the miss rate. Accuracy, response rate, and miss rate were compared between conditions across the 15 participants using two-tailed Wilcoxon signed-rank tests.

### EEG Acquisition and Preprocessing

EEG was recorded continuously from 64 electrodes arranged in the international 10–10 system, sampled at 250 Hz. Online filtering was performed at 1–60 Hz with a 50-Hz notch filter, and signals were referenced to bilateral mastoids; this reference was maintained for all subsequent analyses.

Preprocessing was performed in EEGLAB (Delorme and Makeig, 2004). Artifact Subspace Reconstruction (ASR) was applied to attenuate transient high-amplitude artifacts in the continuous signal. Channels with persistently poor signal quality were identified by visual inspection and removed without interpolation. Independent component analysis (ICA) was then used to identify and remove components reflecting stereotyped artifacts, including blinks, eye movements, and muscle activity, based on scalp topography and time course.

Continuous data were epoched from −720 to 1020 ms relative to object onset (1740-ms epochs). Baseline correction subtracted the mean voltage of the −720 to −500 ms (pre-scene) interval from each epoch. Amplitude-thresholded artifact rejection removed approximately 10% of trials on average. All subsequent analyses were carried out within participant and within condition.

### Time-resolved Decoding

Decoding was performed within each participant using a symmetric sliding window of 11 samples (44 ms at 250 Hz), stepped sample-by-sample from −700 to 1000 ms relative to object onset. Within each window, raw voltages from all retained channels and samples served as features. Because each object identity was consistently paired with a specific background, decoding results are summarized for the post-object interval (0–1 s); pre-object decoding is not reported (Chen et al., 2022).

### Within-condition decoding

For each condition (intact, scrambled), pairwise two-class classification was carried out across all 300 unique pairs of the 25 objects. For each pair and each time window, features were dimensionality-reduced using principal component analysis (PCA), retaining components that explained 99% of the training set variance, and classified with linear discriminant analysis (LDA) using 10-fold cross-validation. Class sizes were approximately balanced (∼30 trials per object after artifact rejection); when minor imbalances remained, all available trials per class were used. Accuracies were averaged across folds and across pairs to yield one decoding time course per condition.

### Cross-condition decoding

To assess context-invariant object information, the same pairwise pipeline was repeated with classifiers trained on trials from one condition and tested on trials from the other (intact→scrambled and scrambled→intact). Training and testing trials were strictly disjoint by condition, so no intact and scrambled trials were ever combined within a training set.

### Temporal generalization

To examine the temporal stability of object codes (King and Dehaene, 2014), we extended the within- and cross-condition pipelines to time-by-time decoding. Classifiers were trained at each time point and tested on all other time points, yielding a two-dimensional temporal generalization matrix for each direction (within-intact, within-scrambled, intact→scrambled, scrambled→intact).

### Representational Similarity Analysis (RSA)

#### EEG RDMs

Time-resolved representational dissimilarity matrices (RDMs) were computed separately for each participant and condition. For each of 100 iterations of a resampling procedure, 30 trials per object were drawn (with replacement when fewer were available), randomly permuted, and split into five batches of six trials. Within each batch, trials were averaged to yield one multichannel response pattern per object at each time point. For each batch and time point, a 25 × 25 RDM was computed by correlating object-evoked patterns across channels using Pearson’s ρ, with dissimilarity defined as 1 − ρ and the diagonal set to zero. The 5 × 100 = 500 RDM estimates per time point were then averaged to yield a single time-resolved RDM per condition and participant (Nili et al., 2014).

#### Model RDMs

We compared neural representational geometry to that of three deep neural networks differing in supervisory regime: ViT large model trained with supervised classification on ImageNet (Dosovitskiy et al., 2020), DINOv2 trained with self-supervised learning (Oquab et al., 2023), and MetaCLIP trained with image–text contrastive learning (Xu et al., 2023). Stimulus images were center-cropped to 250 × 250 pixels (ensuring the target object was contained within the crop) and resized to 224 × 224. To match the grayscale stimuli used in the EEG experiment, images were converted to grayscale and replicated across three channels before model-specific normalization. For ViT and DINOv2, we extracted the last-layer global image embedding; for MetaCLIP, we used the model’s image-feature function. Feature vectors were ordered to match the EEG analyses, and 25 × 25 RDMs were computed by Spearman correlation across feature vectors (1 − ρ; zeros on the diagonal).

#### Model-to-brain alignment

For each participant and condition, each time-resolved EEG RDM was correlated with each model RDM using Spearman’s ρ across upper-triangular off-diagonal entries (Kriegeskorte et al., 2008). Correlations were converted to explained variance (R² = ρ²) and plotted as a function of time.

### Cross-temporal representational similarity analysis

To assess transformations of representational geometry across the scene-preview and post-object periods, we computed cross-temporal similarity between EEG RDMs from the pre-object window (−300 to −50 ms relative to object onset) and the post-object window (0 to 250 ms), separately for intact and scrambled conditions. For each pre- /post-object time-point pair, the upper-triangular entries of the corresponding 25 × 25 RDMs were vectorized and correlated using Spearman’s ρ, yielding a two-dimensional similarity matrix per participant and condition that was then averaged across participants for visualization.

### Model classification paradigm

For each model, a support-vector machine (SVM) classifier was trained on grayscale renders of the training objects to predict the five semantic categories. Across ten splits, the classifier was fit on a random 80% of the training images and tested on the fixed set of 25 objects rendered under intact and scrambled conditions; the same per-split classifier scored both conditions.

### Statistical Analysis

All inferential tests were performed across participants. Behavioral measures like categorization accuracy and response rate were compared between conditions across participants with two-tailed Wilcoxon signed-rank tests. For one-sample tests of time courses against a reference value (e.g., decoding accuracy against the 50% chance level for pairwise classification, or RDM correlations against zero), we used right-tailed Wilcoxon signed-rank tests at each time point. For comparisons between two time courses (e.g., intact vs. scrambled), we used two-tailed paired Wilcoxon signed-rank tests at each time point. For one-dimensional time courses, p-values were corrected for multiple comparisons across time points using the Benjamini–Hochberg false discovery rate (FDR) procedure, applied separately within each analysis and condition (Benjamini and Hochberg, 1995). For two-dimensional maps (temporal generalization matrices, cross-temporal RSA, time-resolved EEG–model alignment), significance was assessed with cluster-based permutation testing (cluster-defining threshold p < 0.05; 1000 permutations).

For each participant, condition, and analysis, the time course was smoothed with a 5-sample moving average, and its peak amplitude was taken as the maximum within the post-object window. Onset latency was defined as the earliest post-object time point at which the time course first reached, and then remained at or above, 60% of this peak for at least 12 ms. The criterion was computed relative to the reference level as reference + 0.60 × (peak − reference), with the reference set to 50% for decoding accuracy and to 0 for R² and for between-condition difference time courses. Peak latency was the time of the peak and peak amplitude its value. These three measures were computed identically across the within-condition decoding, cross-condition decoding, and model-to-brain RSA analyses, estimated per participant, and compared between conditions with two-tailed Wilcoxon signed-rank tests; values are reported as mean ± SEM. The smoothing affected only these latency and amplitude measures, not the statistical tests.

DNN model’s classification accuracy was compared between conditions with a per-object paired Wilcoxon signed-rank test across the three models.

## Code Accessibility

Code for this study is publicly available on GitHub, and will be released upon publication of the paper.

## Data Availability

The EEG data and stimulus materials that support the findings of this study are available from the corresponding author upon reasonable request.

## Acknowledgments

This research received no specific grant from any funding agency in the public, commercial, or not-for-profit sectors. No generative AI or AI-assisted tools were used in the generation, analysis, or visualization of data, including figures. AI-assisted tools were used solely for language editing to improve clarity and readability.

