## Supplementary Materials for "Coherent scene context accelerates and reshapes neural object representations"

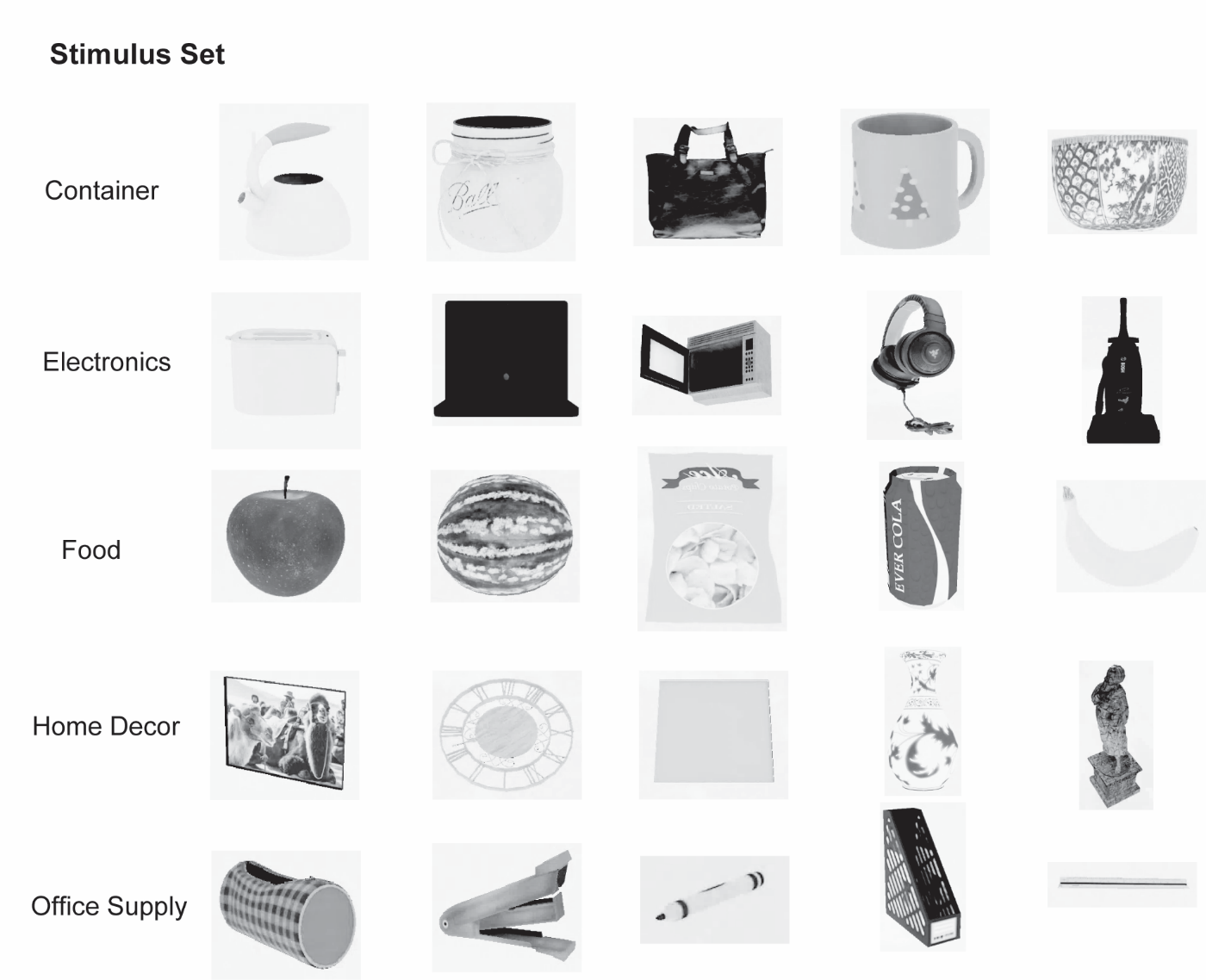

**Figure S1. Complete stimulus set.**

*All 25 object identities, five per semantic category (containers, electronics, food, home décor, and office supplies). All images were presented in grayscale to remove color-based saliency.*

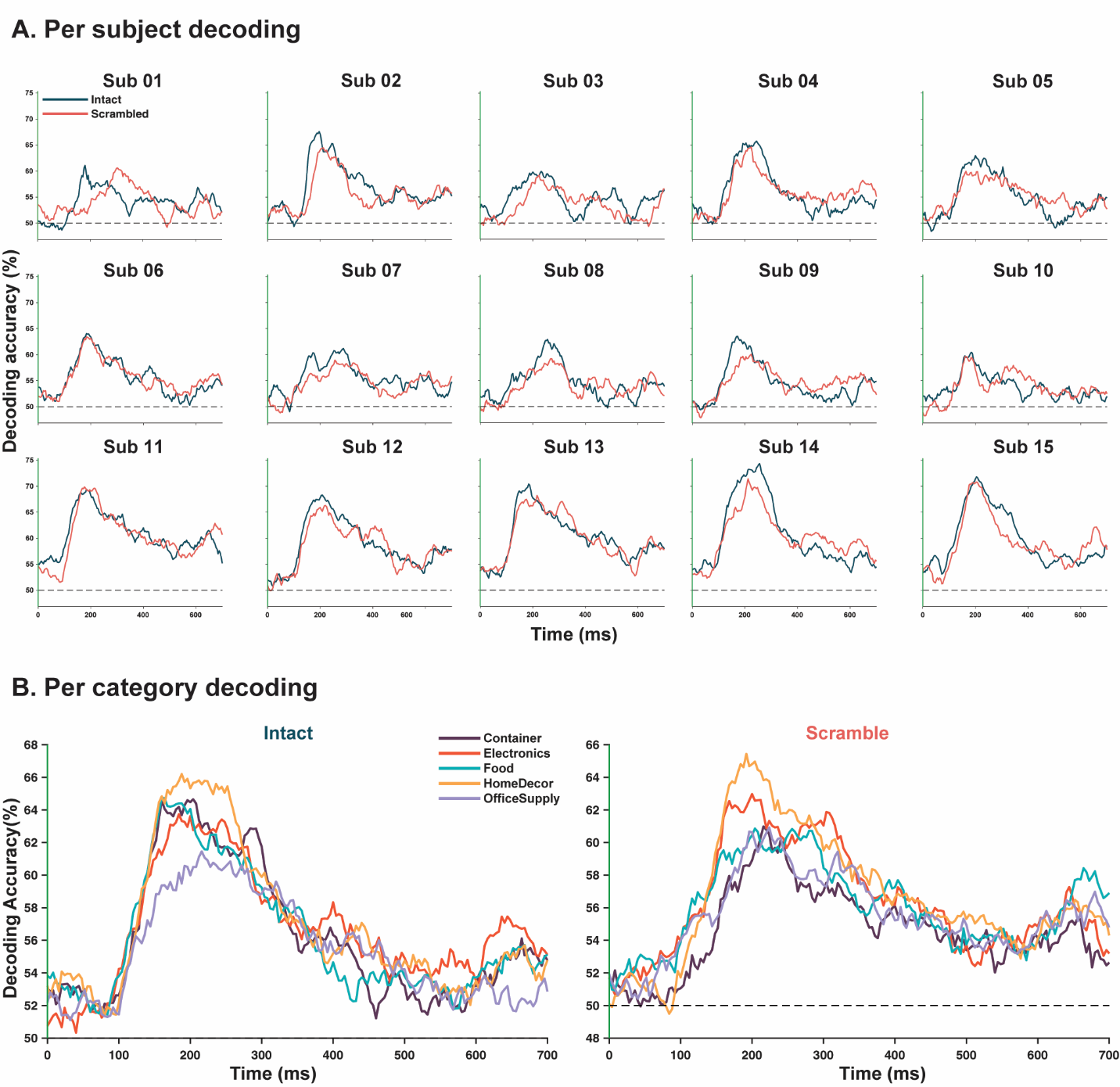
**Figure S2. Per-subject and per-category object decoding.**

**A.** *Time-resolved pairwise decoding for each of the 15 participants, intact (blue) and scrambled (pink). Dashed lines indicate chance (50%).*

**B.** *Time-resolved decoding averaged within each semantic category (containers, electronics, food, home décor, office supplies) for the intact and scrambled conditions. The intact-over-scrambled advantage and the early decoding rise were consistent across participants and categories.*

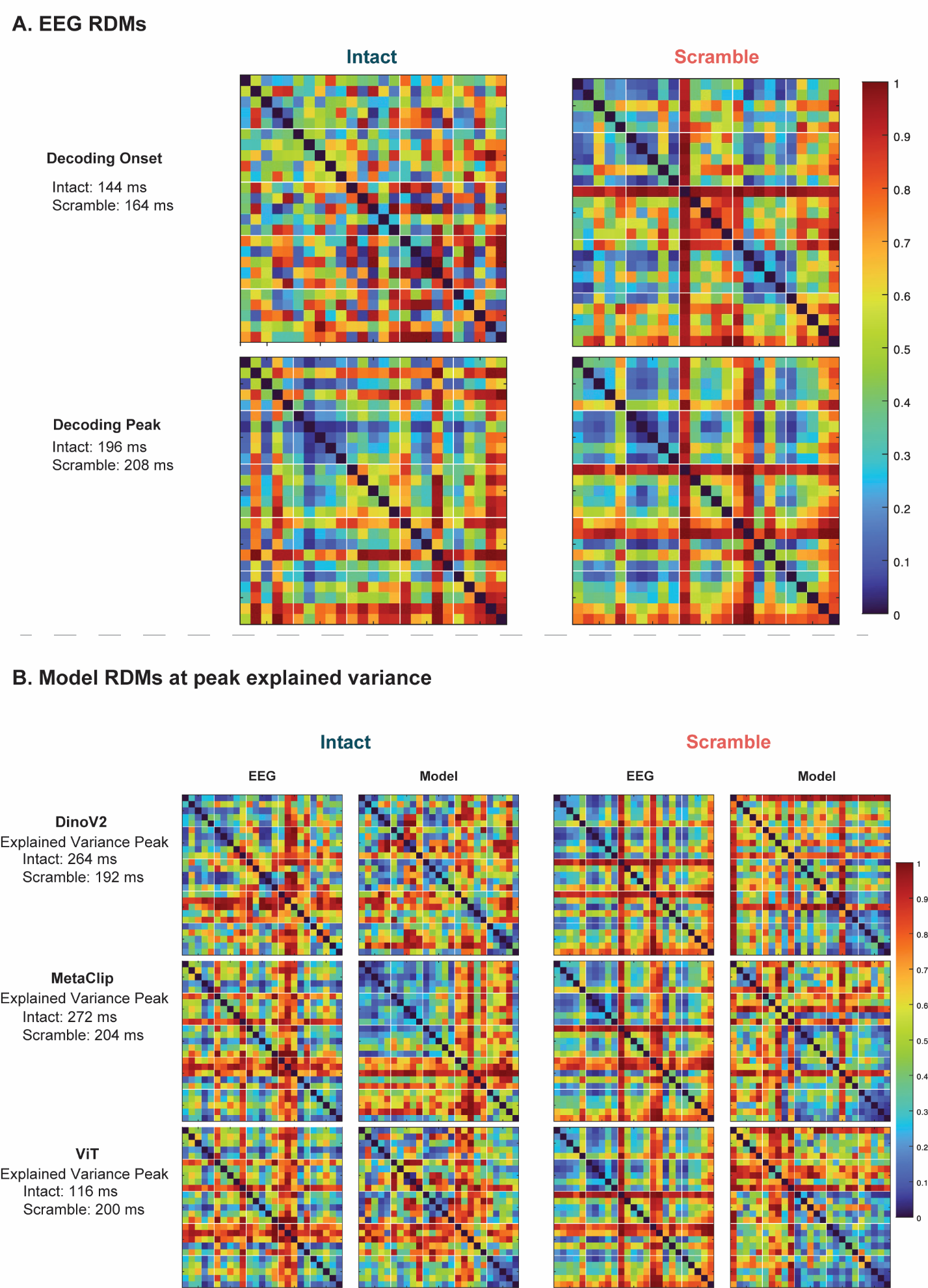

**Figure S3. EEG and model representational dissimilarity matrices.**

**A.** *Group-averaged 25 × 25 EEG RDMs for the intact and scrambled conditions at decoding onset (intact 144 ms, scramble 164 ms) and decoding peak (intact 196 ms, scramble 208 ms).*

**B.** *EEG and model RDMs at each model's peak explained variance for DINOv2 (intact 264 ms, scramble 192 ms), MetaCLIP (intact 272 ms, scramble 204 ms), and ViT (intact 116 ms, scramble 200 ms). Color indicates dissimilarity (1 − Spearman's ρ). MDS embeddings of these same matrices are shown in Figure S4.*

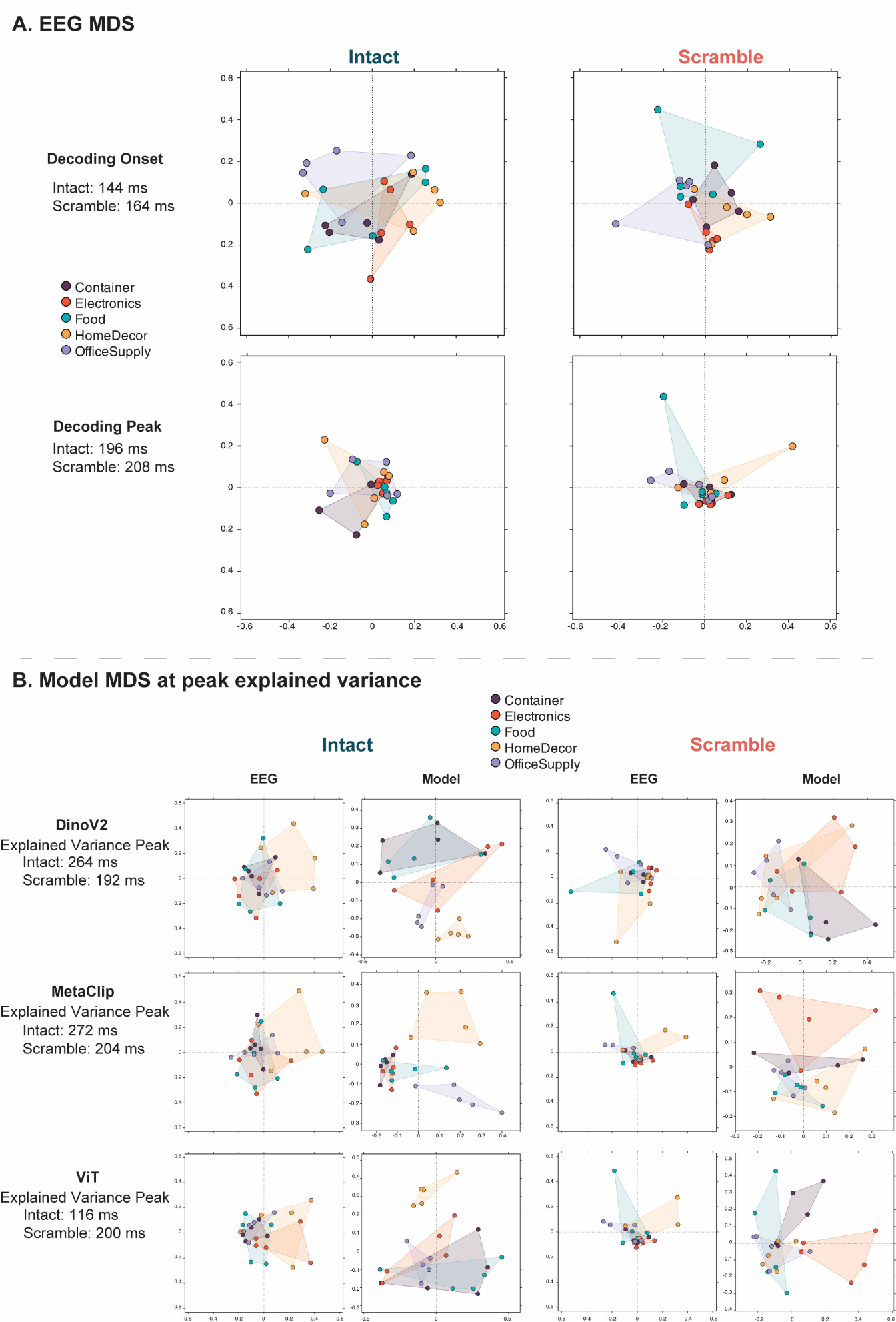

**Figure S4. Multidimensional scaling of EEG and model representational geometry.**

*Two-dimensional MDS embeddings of the RDMs in Figure S3, with objects colored by semantic category.*

**A.** *EEG geometry at decoding onset and decoding peak for the intact and scrambled conditions.* **B.** *EEG and model geometry at each model's peak explained variance for DINOv2, MetaCLIP, and ViT. Shaded hulls connect objects within a category.*

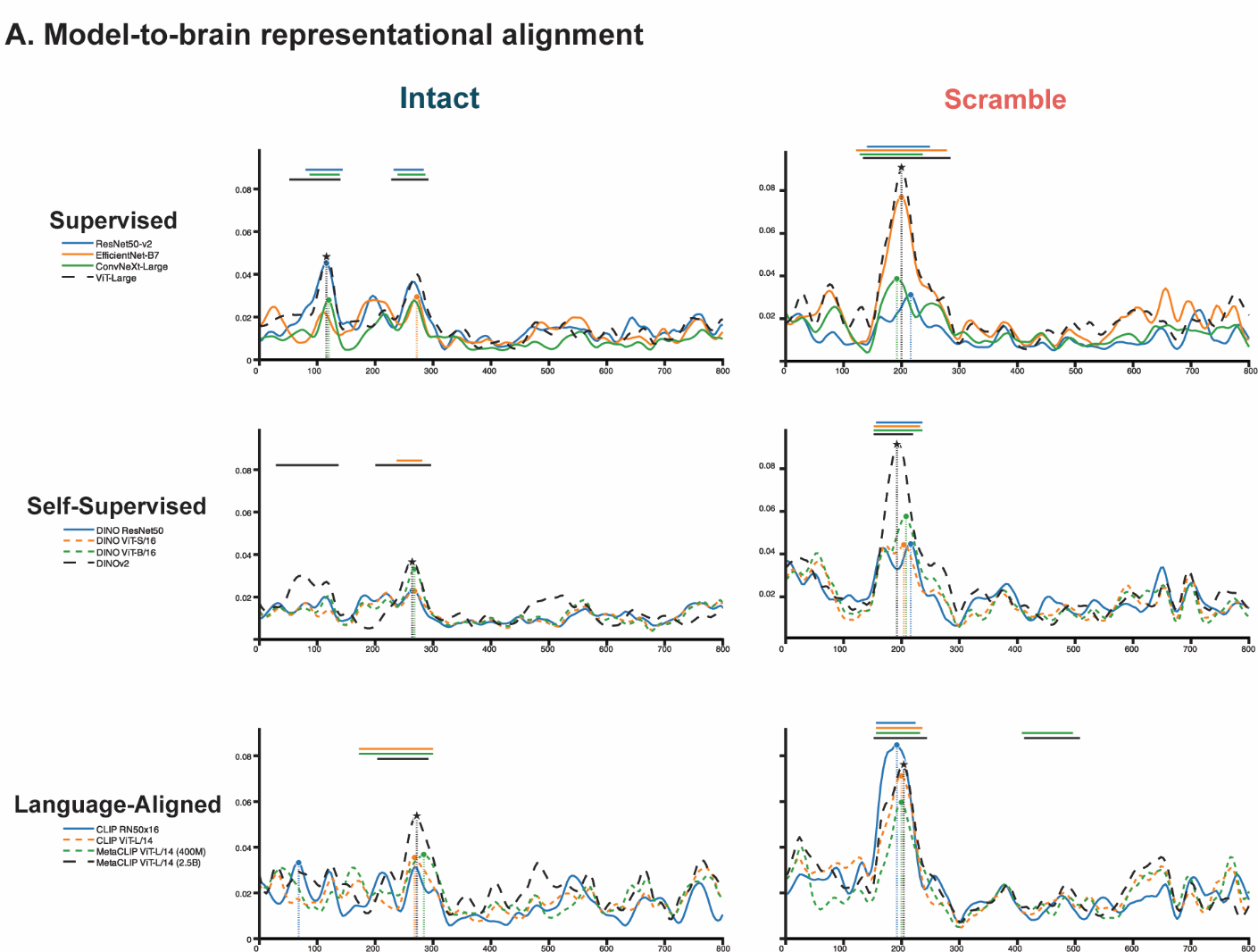

**Figure S5. Model-to-brain representational alignment across architectures.**

*Time-resolved explained variance (R²) between the EEG RDM and each model RDM in the intact and scrambled conditions, shown for three supervisory regimes: supervised vision (ResNet50-v2, EfficientNet-B7, ConvNeXt-Large, ViT-Large), self-supervised vision (DINO ResNet50, DINO ViT-S/16, DINO ViT-B/16, DINOv2), and language-aligned vision–language models (CLIP RN50x16, CLIP ViT-L/14, MetaCLIP ViT-L/14 [400M], MetaCLIP ViT-L/14 [2.5B]). Shaded regions denote ±1 SEM across participants. Horizontal bars indicate time points of significant model–brain alignment for each model (cluster-corrected, p < 0.05). Stars mark peak alignment for the model used in the result section, circles mark peak alignment for the models used to check consistency, and dotted vertical lines mark peak latency. The late, intact-specific alignment advantage of language-aligned models generalized across architectures within each family.*
